# Social Isolation Intensifies *adgrl3.1*-Related Externalizing and Internalizing Behaviors in Zebrafish

**DOI:** 10.1101/2024.09.19.613974

**Authors:** Barbara D. Fontana, Nancy Alnassar, William H.J. Norton, Matthew O. Parker

## Abstract

Externalizing disorders (EDs) are characterized by outward-directed behaviors such as aggression and hyperactivity. They are influenced by gene-environment interactions, yet our understanding of the genetic predispositions and environmental contexts that give rise to them is incomplete. Additionally, people with EDs often exhibit comorbid internalizing symptoms, which can complicate the clinical presentation and treatment strategies. Following on from our previous studies, we examined genes x environment interaction as a risk factor for EDs by looking at internalizing and externalizing behaviors after social isolation. Specifically, we subjected *adgrl3*.*1* knockout zebrafish — characterized by hyperactivity and impulsivity — to a 2-week social isolation protocol. We subsequently assessed the impact on anxiety-like behavior, abnormal repetitive behaviors, working memory, and social interactions. Genotype-specific additive effects emerged, with socially isolated *adgrl3*.*1* knockout fish exhibiting intensified comorbid phenotypes, including increased anxiety, abnormal repetitive behaviors, reduced working memory, and altered shoaling, when compared to WT fish. The findings demonstrate that genetic predispositions interact with environmental stressors, such as social isolation, to exacerbate both externalizing and internalizing symptoms. This underlines the necessity for comprehensive diagnostic and intervention strategies.

## 1. Introduction

Externalizing disorders (EDs) encompass a range of conditions characterized by outward-directed behaviors such as aggression, rule-breaking, impulsivity, and hyperactivity (Liu, 2004). These disorders, including conduct disorder, oppositional defiant disorder, and substance abuse disorders, pose significant challenges for individuals and public health systems (de Lacy and Ramshaw, 2023). The high heritability of EDs underscores the importance of understanding the genetic and environmental factors that contribute to these behaviors (Samek and Hicks, 2014). One of the genes frequently associated with EDs is the adhesion G-protein-coupled receptor L3 (*ADGRL3*; also known as *LPHN3*) gene. This gene is linked to an increased risk of several externalizing disorders, including ADHD, in humans (Domene et al., 2011; Hwang et al., 2015). ADGRL3 plays a crucial role in cell adhesion, signal transduction, and synaptic signaling (Arcos-Burgos et al., 2010; McMillan et al., 2002; Scholz et al., 2015). In zebrafish, the behavioral and molecular mechanisms underlying *adgrl3*.*1*-related ED phenotypes have been studied in larvae (Lange et al., 2018; Lange et al., 2012; Sveinsdóttir et al., 2023) and, more recently, in adult animals (Fontana et al., 2023). Adult *adgrl3*.*1* knockout zebrafish exhibit behaviors such as hyperactivity, impulsivity, and attention deficits, which are core features of EDs (Fontana et al., 2023).

An important factor affecting the quality of life of patients with externalizing disorders is their social interactions and inclusion in society (Meßler et al., 2016; Shaw-Zirt et al., 2005). In addition, internalizing symptoms often co-occur in patients with EDs, and can be attributed to socio-developmental factors (Ray et al., 2017; Speyer et al., 2021). Internalizing symptoms are psychological issues that manifest ‘internally’ within an individual, such as anxiety and social withdrawal, causing distress that is not immediately visible to others (Lilienfeld, 2003). Genetic predispositions to externalizing behavioral patterns give rise to challenging family, educational and peer dynamics, leading to social exclusion or isolation and thus exacerbating internalizing symptoms (Kwan et al., 2020; Smit et al., 2020). For example, patients with ADHD often experience social difficulties and rejection due to their inattention, impulsivity and hyperactivity, and rejection is associated with increased aggression and lower IQ in ADHD children (Carpenter Rich et al., 2009). The rejection and social exclusion then lead to a vicious cycle, where these patients’ lack of social contact or rejection will be highly associated with other comorbidities increasing the severity of symptoms, and making it more difficult to treat these disorders (Hoza, 2007). Internalizing disorders are often comorbid in ADHD children who report having fewer friendships, more negative friendship quality and loneliness (Smit et al., 2020), raising the importance of understanding social isolation in EDs and its comorbidities. Social exclusion or isolation may not only affect the severity of symptoms but also cause negative effects on immunity and stress-related pathways across species (Forsatkar et al., 2017; Ieraci et al., 2016; Tuchscherer et al., 2004; Zlatković et al., 2014). Understanding these underlying reasons can aid in better diagnosis and targeted interventions.

Previously we showed an overlap of genes linked to the behavioral phenotypes found in *adgrl3*.*1* knockouts and socially isolated WT animals (Alnassar et al., 2023; Fontana et al., 2023) supporting the idea that social isolation could result in a worsening of EDs and its comorbidities, such as anxiety. Here, we aimed to explore the effects of social isolation on a well-characterized model of EDs by using zebrafish that lack *adgrl3*.*1* gene function. Understanding how social isolation impacts *adgrl3*.*1*^*-/-*^ zebrafish is essential to elucidate the environmental modulation of genetic risk factors and providing a broader view of how social factors influence the neurobiology of externalizing behaviors. We hypothesized that social isolation would significantly exacerbate behavioral phenotype severity, highlighting the crucial role of the environment in the interaction between ADGRL3 function and the etiology of EDs. To test this hypothesis, zebrafish were isolated for 14 days without any visual or olfactory contact with conspecifics and subsequently assessed using various behavioral tasks, including the shoal test, novel tank diving test, novel object boldness task, and FMP Y-maze. Our findings reveal, for the first time, the baseline behavior of *adgrl3*.*1* knockout zebrafish compared to wild-type (WT) in a social behavior task, providing valuable insights into the interplay between genetic and environmental factors in externalizing disorders.

## 2. Material and Methods

### 2.1. Animal husbandry and experimental design

The *adgrl3.1*^*-/-*^ zebrafish line was generated by CRISPR-Cas9 technology as previously described (Sveinsdóttir et al., 2023) and compared with age-matched wild-type (WT) zebrafish (AB strain) (total *n* = 128; 50:50 male: female ratio). Animals were bred in-house and fed with flake food (ZM-flake and ZM-300, ZM Ltd.) three times a day and adult brine shrimp during the mornings. The animals were grown in a re-circulating system on a 14/10-hour light/dark cycle (lights on at 9:00 a.m.), pH 8.4, at 28 °C (±1 °C). Zebrafish were kept in groups of 10 animals per 2.8 L until 4 months post fertilization (adult age) when they were divided into four groups (WT group-housed; WT social isolated; *adgrl3.1* group-housed; *adgrl3.1* social isolated). Group housed fish were grown and kept in 2.8 L tanks with 10 animals per tank until the experiment day when they were behaviorally tested. Meanwhile, individually housed fish were kept in smaller tanks (1 L of water) with no visual or olfactory cues to their conspecifics for two weeks before being behaviorally tested (**Fig. 1**).

**Fig. 1.**
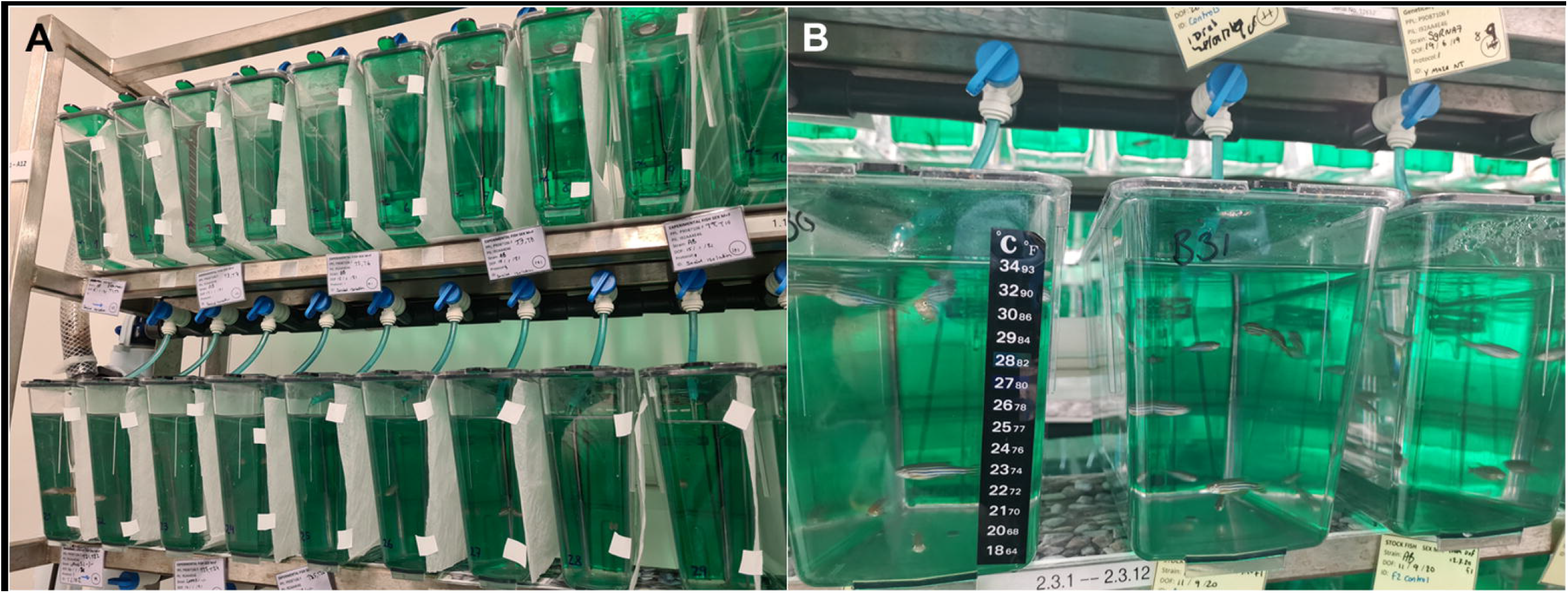
Picture of the experimental setup used for social isolation. **(A)** Shows the individual housing tanks, where animals had no shared water or visual contact with conspecifics. **(B)** Housing tanks used to keep fish in groups in a density of 2 fish/L and with visual contact to other fish tanks.

On the first experimental day, fish were tested in the novel tank diving test (NTT) (*n* =16) or novel object boldness task (*n* = 16). After, fish were immediately transferred to the FMP Y-maze (*n* = 20) or and then returned to their previous social context. 24 h later, fish were tested in the shoal test (*n* of shoal *=* 8; 32 animals per group) and then culled. Animals were tested at the shoal test on the second day of the experiment to avoid any kind of social interactions prior to the assessment of other behavioral domains. The number of animals per group were calculated a priori (d = 0.4, power = 0.75, alpha = 0.05) following extensive published work from our laboratory using zebrafish as a translational model (Alnassar et al., 2023; Fontana et al., 2020; Fontana et al., 2023) and considering our primary outcomes which resulted in a sample size of 66 animals. For the shoal test, sample size was decided based on previous publications considering that a *n* of 8 shoals uses 32 animals per group. Once the experiment was over fish were euthanized using 2-phenoxyethanol from Aqua-Sed (Aqua-Sed™, Vetark, Winchester, UK). Extreme outliers were removed from analysis [>3*IQR], being four animals for NTT, one animal from FMP Y-maze and one shoal from social behavior test. Researchers were blind to the experimental conditions (fish genotype) to avoid biased handling during the behavioral sessions and animals were randomly selected from one of four tanks for each experimental design. All behavioral tests were performed between 09:00 a.m. and 03:00 p.m. and were approved by the Animal Welfare and Ethical Review Board, and under license from the UK Home Office (Animals (Scientific Procedures) Act, 1986) [PPL: P9D87106F].

### 2.3. Novel tank diving test (NTT)

The NTT is used to assess locomotor and anxiety-related phenotypes in zebrafish (Egan et al., 2009; Kalueff et al., 2013; Levin et al., 2007; Wong et al., 2010). Group housed *vs*. isolated animals were tested for both the WT and *adgrl3.1* genotype and they were individually placed in the novel tank (20 cm x 17.5 cm x 5 cm; L x H x W) containing 1 L of aquarium water for 6 minutes (Egan et al., 2009; Parker et al., 2012; Rosemberg et al., 2012). Behavioral activity was analyzed using the Zantiks automatic system (Zantiks Ltd., Cambridge, UK) and the analysis parameters such as time spent in bottom (tank divided in three virtual areas) and distance traveled were considered to evaluate anxiety and locomotion, respectively. Animals were returned to their housing system after the novel tank diving test and tested on the shoal behavior test 24 hours later.

### 2.4. Novel object boldness task

The novel object boldness task was previously used by different groups (Norton et al., 2011; Wright et al., 2003) and assesses an animal’s willingness to approach a novel object that can be perceived as a threat (15-cm falcon tube made with black clay). This boldness behavior can also be defined as individuals’ disposition to take risks and is also often linked to risk-taking behavior (Sih et al., 2004). Here, condition (isolation or not) and genotype (*adgrl3.1 vs*. WT) were considered when assessing the time that animals spent close to the object (2 cm distance around the object). Animals were recorded for 6 minutes with a webcam and videos further analyzed using ANY-maze behavioural tracking software. The tank dimensions were 27 cm x 36 cm x 10 cm (W x L x D) and animals were submitted to the FMP Y-maze after this behavioral protocol.

### 2.5. FMP Y-maze

The FMP Y-maze task is a cognitive task validated for measuring zebrafish working memory and cognitive flexibility (Cleal et al., 2021; Fontana et al., 2019) and has been recently shown that it has important relevance for assessing abnormal repetitive behavior in this same species (Fontana et al., 2021). Zebrafish were used for assessing FMP Y-maze performance in a white Y-maze tank with three identical arms (5 cm length x 2 cm width; 120º angle between arms) filled with 3L of aquarium water. The tank had no explicit intra-maze cues and was lit with the ambient light of 1 LUX to allow some visibility in the maze. Behavior was recorded for 60 minutes using the Zantiks automatic AD system (Zantiks Ltd., Cambridge, UK) and analysis was made based on overlapping series of four choices (tetragrams) (Gross et al., 2011). Working memory and repetitive behavior were assessed using the number of alternations (rlrl + lrlr) and repetitions (rrrr + llll) as a proportion of the total number of turns which are highly expressed through 1-hour (% of total turns) (Cleal and Parker, 2018; Gross et al., 2011). Moreover, specific alterations and repetitions across time (10 min time bins) were analyzed in-depth to assess cognitive flexibility. After the FMP Y-maze task fish were also returned to their original housing system for further analysis of shoal behavior.

### 2.6. Shoal behavior

24 h later after novel tank diving task or FMP Y-maze task, fish (4-fish per shoal) were simultaneously placed in the test tank (25×15×10 cm length x height x width) and group behavior was analyzed for 5 minutes (Canzian et al., 2017; Green et al., 2012; Muller et al., 2017; Schmidel et al., 2014). The Zantiks automatic AD unit (Zantiks Ltd., Cambridge, UK) was used to record the fish behavior, and the distance between fish and shoal area was analyzed using 15s screenshots during 5 min trials (20 screenshots *per* trial) in the Image J 1.49 software (Green et al., 2012; Schmidel et al., 2014). All screenshots were calibrated proportionally to the size of the tank.

### 2.8. Statistics

Data from the novel tank diving test, open field test, FMP Y-maze and shoal behavior was analyzed using two-way ANOVA with genotype (two levels – WT *vs. adgrl3.1*) and social context (two levels – group house *vs*. social isolation) as fixed factors. Two-way RM-ANOVA was used to analyze data across time in the FMP Y-maze using time (six levels – 10, 20, 30-, 40-, 50- and 60-time bins) and group (four levels – WT, WT + social isolation, *adgrl3.1*^*-/-*^ and *adgrl3.1*^*-/-*^ + social isolation) as main factors. Tukey’s test was used as a post-hoc analysis, data were represented as mean and error of the mean (SEM), and results were considered significant when p ≤ 0.05.

## 3. Results

### 3.1. Social isolation differently affects anxiety-like behavior depending on genotype

First, we investigated anxiety-related phenotypes and risk-taking behaviors of WT and *adgrl3.1*^*-/-*^ zebrafish, with or without social isolation (**Fig. 2)**. No isolation (F _(1, 56)_ = 0.4072; *p* = 0.5260) or interaction effect (isolation *vs*. genotype; F _(1, 56)_ = 1.917; *p* = 0.1716) were observed in animals’ locomotion (distance traveled), however a genotype effect (F _(1, 56)_ = 7.205; *p\*\** = 0.0095) was observed where *adgrl3.1*^*-/-*^ animals had a decreased distance traveled compared to WT (*p\** = 0.0169). Genotype had also a significant effect on time spent immobile (F _(1, 56)_ = 31.68; *p**** <* 0.0001) where both *adgrl3.1*^-/-^ (*p\*\*\** = 0.0003) and *adgrl3.1*^-/-^ + social isolation (*p\*\*\** = 0.0006) spent more time immobile compared to controls. Although no social isolation effect was observed for time spent in bottom (F _(1, 56)_ = 1.543; *p* = 0.2194), both genotype (F _(1, 56)_ = 84.40; *p**** <* 0.0001) and interaction between factors (F _(1, 56)_ = 8.754; *p\*\** = 0.0045) was observed for this parameter. Post-hoc analysis yielded a significant increase in the time spent bottom for *adgrl3.1*^-/-^ (*p\*\*\** = 0.0003) and *adgrl3.1*^-/-^ + social isolation (*p**** <* 0.0001) compared to controls. Moreover, WT social isolated animals showed a significant decrease in the time spent in bottom compared to WT group housed (*p\** = 0.0194) (**Fig.2A**).

**Fig. 2.**
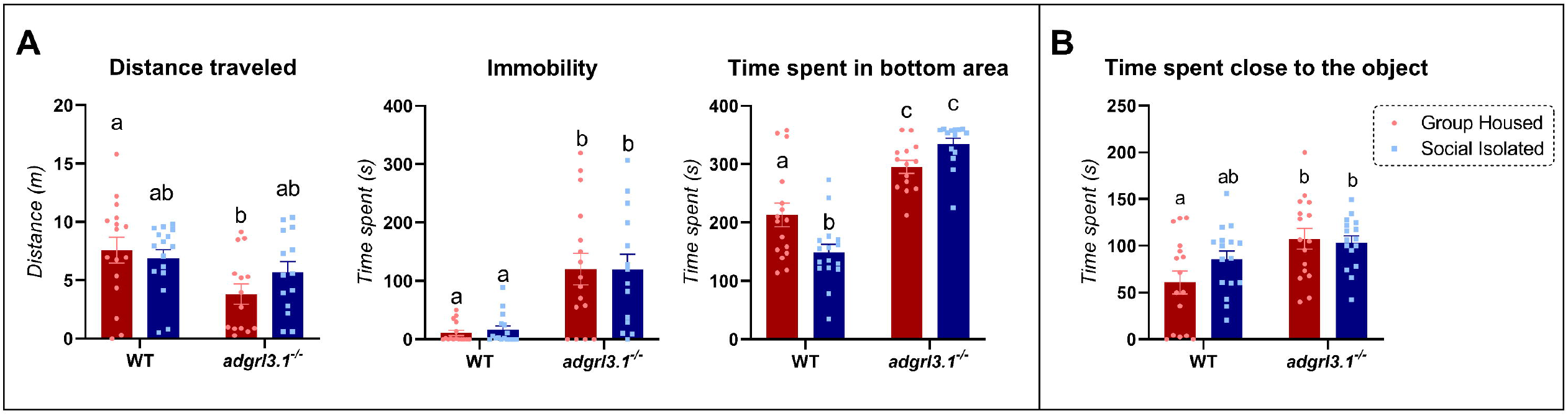
Effects of social isolation on **(A**) locomotor and anxiety-related parameters (*n* = 14 – 16) and **(B)** risk-taking behavior (*n* = 16). Data were represented as mean ± S.E.M and analyzed by two-way ANOVA (social isolation *vs*. genotype), followed by Tukey’s test multiple comparison test between groups. Different letters indicate statistical differences between groups (*p* < 0.05).

For risk-taking behavior (**Fig. 2B**), only a significant effect of genotype was found (F _(1, 60)_ = 10.52; *p\*\** = 0.0019) with neither effect for social isolation (F _(1, 60)_ = 1.077; *p* = 0.3035) nor an interaction effect between both factors (F _(1, 60)_ = 2.001; *p* = 0.1624).

### 3.2. Working memory and cognitive flexibility of adgrl3.1^-/-^ zebrafish after social isolation

Following two-way ANOVA, social isolation (F _(1, 75)_ = 17.78; *p**** <* 0.0001) had a main effect on exploratory activity of the animals when looking at average turns, with no genotype (F _(1, 75)_ = 0.3177; *p* = 0.5746) or interaction effect (F _(1, 75)_ = 2.080; *p* = 0.1534). A decrease in the average turns were observed for isolated WT compared to their own group housed controls (*p\*\*\** = 0.0010). When looking at alternations and repetitions, no social isolation effect (F _(1, 75)_ = 1.908; *p* = 0.9483 and F _(1, 75)_ = 2.404; *p* = 0.1253) nor interaction between factors (F _(1,75)_ = 0.004; *p* = 0.9483 and F _(1, 75)_ = 0.007; *p* = 0.9331) was observed. However, a genotype effect was observed in both alternations (F _(1,75)_ = 21.02; *p**** <* 0.0001) and repetitions (F _(1,75)_ = 16.76; *p**** <* 0.0001). Briefly, alternations were only decreased in *adgrl3.1* knockout and social isolated (*p\** = 0.0263) compared to WT group housed. Meanwhile, for repetitions, both mutant groups group housed (*p\** = 0.0417) and social isolated (*p\*\** = 0.0023) had a significant increase in repetitions compared to WT group housed (**Fig. 3A**).

**Fig. 3.**
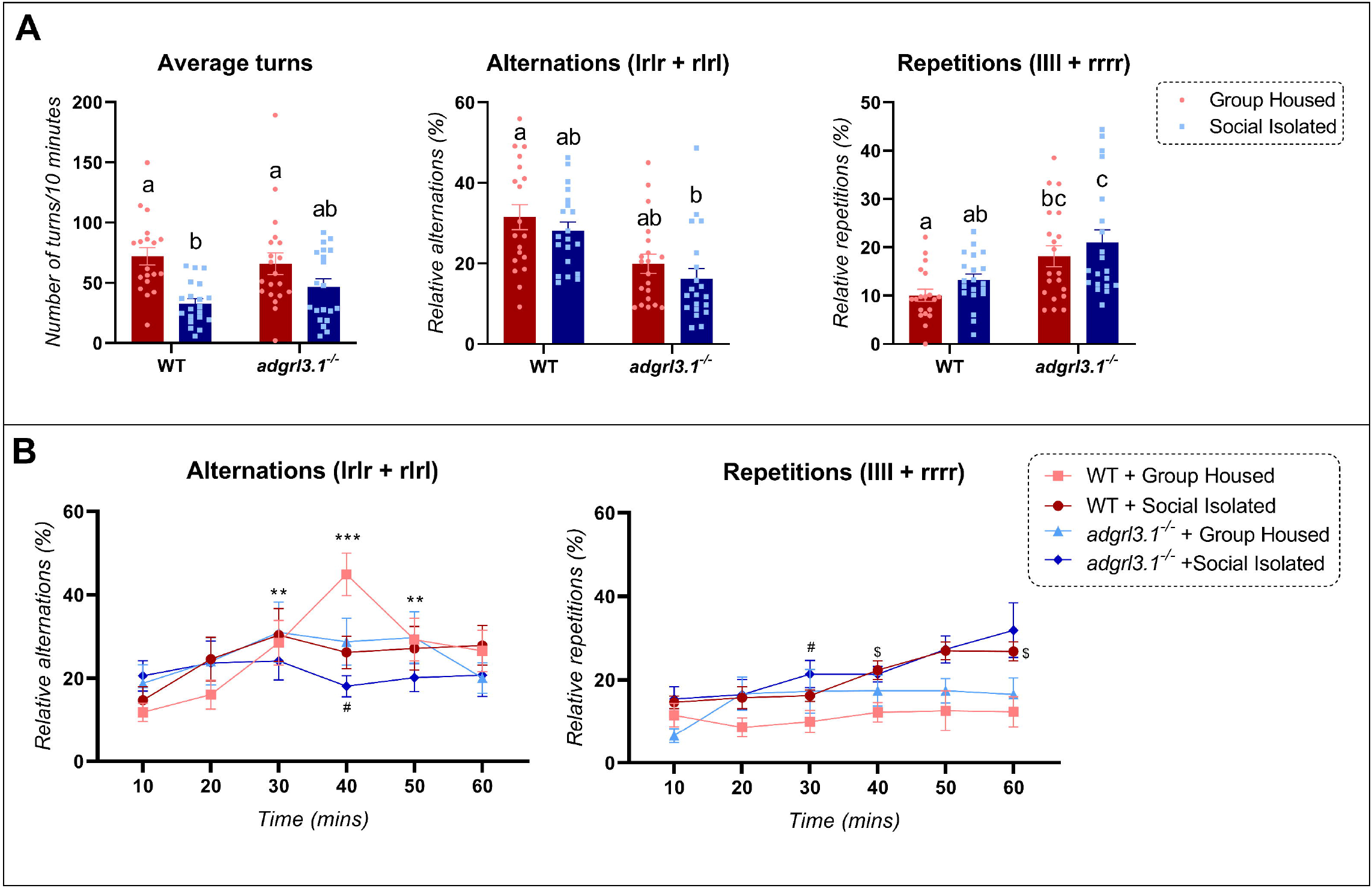
Social isolation effects on the FMP Y-maze. **(A)** Zebrafish knockout for *adgrl3.1* gene show decreased alternation when socially isolated. **(B)** Alterations in cognitive flexibility are observed in all social isolated and *adgrl3.1* group, meanwhile, repetitions are significantly higher at 60-time bin for *adgrl3.1*^*-/-*^ + social isolation compared to WT group-housed. Data were represented as mean ± S.E.M and analyzed by two-way ANOVA (social isolation *vs*. genotype or time *vs*. group), followed by Tukey’s test multiple comparison test between groups. Different letters indicate statistical differences between groups (*p* < 0.05), asterisks indicate significant difference compared to 10 min time bin (*p\** < 0.05), octothorp means significant difference compared to WT group-housed for the same time bin (*p*^#^ < 0.05) and $ mean significant difference of WT compared to WT social isolated for the same time bin (*p*^*$*^ < 0.5; *n* = 19 - 20).

**Fig. 3B** shows the effects of social isolation on WT and knockout animals for the *adgrl3.1* gene when looking at cognitive flexibility analysis. Although no effect was observed for the different groups (F _(3, 75)_ = 0.4426; *p* = 0.7233) a significant time effect was observed for alternations across time (F _(3.984, 298.8)_ = 6.389; *p**** <* 0.0001) and interaction between factors (F _(15, 375)_ = 1.466; *p\*\** = 0.0030). Only the WT group housed had an increase on alternations at 30,40 and 50 min compared to 10 min time bin (*p* <* 0.05). At 40 min, peak of alternations, the group *adgrl3.1* + social isolation showed a decrease in alternations compared to group housed WT.

Two-way RM-ANOVA analysis of repetitions across time yielded a significant effect for time (F _(3.723, 279.2)_ = 8.262; *p**** <* 0.0001) and interaction time*group (F _(3, 75)_ = 5.088; *p**** <* 0.0001), with no interaction effect F _(15, 375)_ = 1.592; *p* = 0.0732). WT zebrafish social isolated had an increase in repetitions at 40- and 60-min time bin compared to WT group housed (*p* <* 0.05), meanwhile social isolated knockout animals showed an increase at repetitions only at 30 min (*p\** < 0.05).

### 3.3. adgrl3.1 knockout animal show disrupted shoaling behavior

Shoaling behavior was analyzed for 5 min and the effects of social isolation in animals’ knockout for the *adgrl3.1* gene can be observed in **Fig. 4**. Although no effect was observed for the social isolation *per se* (F _(1, 27)_ = 0.3548; *p* = 0.5564), a genotype (F _(1, 27)_ = 11.94; *p\*\** = 0.0018) and interaction effect between factors (F _(1, 27)_ = 6.471; *p\** = 0.0170) were observed for shoal area. Shoal area was increased for *adgrl3.1*^*-/-*^ group (*p\*\** = 0.0010) with no effect for *adgrl3.1*^*-/-*^ + social isolation (*p* = 0.2171) when comparing these groups to WT group housed. No significant differences were observed for the WT *vs*. WT social isolated. Similarly, a genotype (F _(1, 27)_ = 13.87; *p\*\*\** = 0.0009) and interaction effect (F _(1, 27)_ = 4.367; *p\** = 0.0462) were observed for inter-fish distance with no effect for social isolation (F _(1, 27)_ = 0.2190; *p* = 0.6436). Tukey’s post-hoc analysis showed a significant effect for *adgrl3.1*^*-/-*^ when compared to WT group housed (*p\*\** = 0.0015). No significant differences were observed for both social isolated groups, WT (*p* = 0.6522) and *adgrl3.1*^*-/-*^ (*p* = 0.1320), compared to WT group housed.

**Fig. 4.**
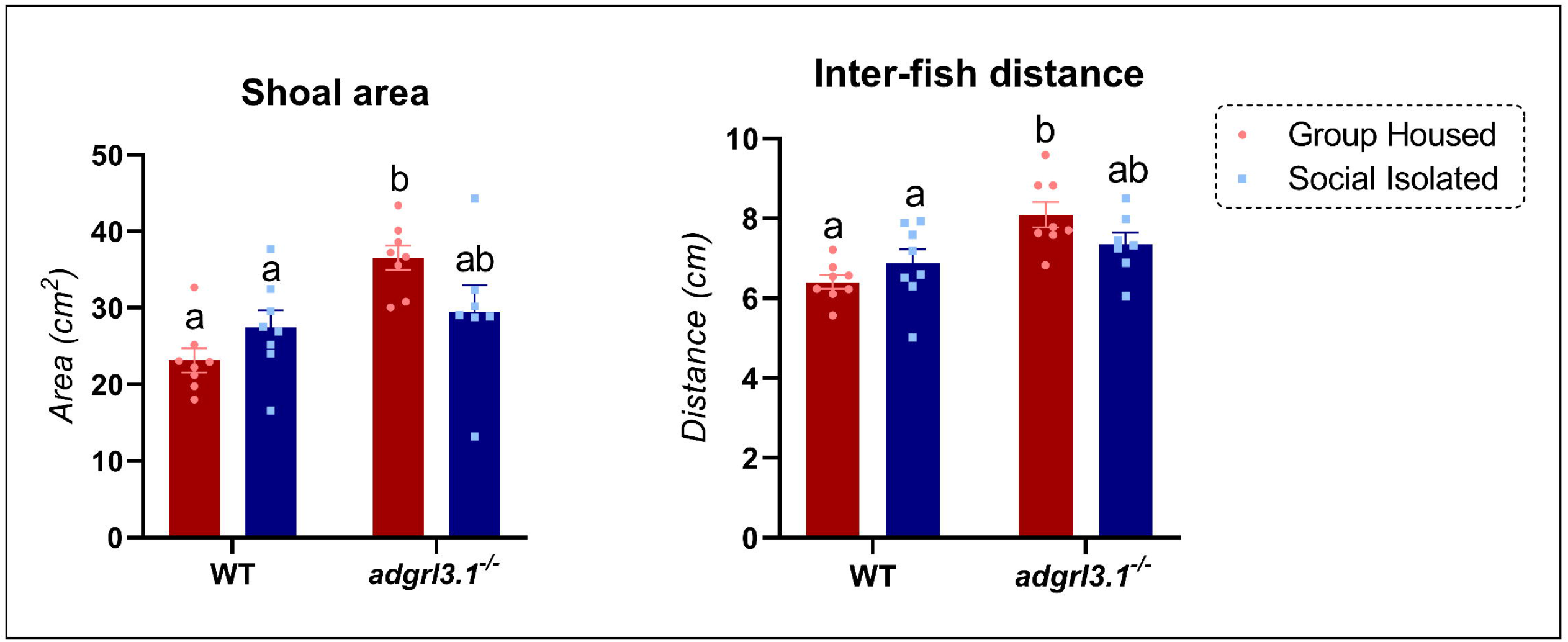
Shoaling behavior of socially isolated animals’ knockout for the *adgrl3.1* gene. Data were represented as mean ± S.E.M and analyzed by two-way ANOVA (genotype *vs*. social isolation), followed by Tukey’s test multiple comparison test between groups. Different letters indicate statistical differences between groups (*p* < 0.05; *n* = 7 – 8).

## 4. Discussion

Motivated by our previous work, which revealed significant overlap between the molecular pathways altered in *adgrl3.1*^*-/-*^ zebrafish and in wild-type zebrafish subjected to social isolation (Alnassar et al., 2023; Fontana et al., 2023), this study investigated the effects of a sustained two-week social isolation period on the behavior of *adgrl3.1*^*-/-*^ zebrafish. Our aim was to test the hypothesis that social isolation would exacerbate abnormal phenotypes observed on those mutants. Social isolation has a strong interaction with the behavioral phenotypes shown by *adgrl3.1*^*-/-*^, with isolated knockout animals showing opposing effects of both internalizing-like and externalizing-like behaviors. For example, in the FMP Y-maze, *adgrl3.1*^*-/-*^ fish showed increases in repetitive behavior and decreases in cognitive flexibility. However, following social isolation, *adgrl3.1* knockout also showed decreased working memory, together with the other behavioral changes observed in the non-isolated animals. Social behavior was also affected in the *adgrl3.1* knockout, with increased shoal area and inter-fish distance observed, suggesting disrupted social behavior in the presence of reduced *adgrl3.1* function. Social isolation also affected the disrupted social behavior in *adgrl3.1* knockout zebrafish by attenuating those effects in terms of shoal area and inter-fish distance. Interestingly, the effects of social isolation in WT such as decreased social behavior and anxiety-like behavior were observed as an opposite trend for *adgrl3.1* knockout, showing that social isolation affects animals differently depending on their genotype.

Social disruption throughout life can alter physiological, neurochemical, neuroendocrine, and behavioral functions (Matthews and Tye, 2019; Mumtaz et al., 2018). These pronounced changes are associated with altered brain function (Fabricius et al., 2010; Lapiz et al., 2003; Schoenfeld and Gould, 2012; Westenbroek et al., 2004) and the activation of the hypothalamic-pituitary-adrenal (HPA) axis (Cacioppo et al., 2000; Serra et al., 2005). In zebrafish, socially isolated animals often show blunted cortisol response to acute stressors (Giacomini et al., 2015) together with altered internalized behaviors such as decreased anxiety-like behavior (Parker et al., 2012). Here, we also observed a decreased time spent in bottom when comparing WT group-housed *vs*. WT social isolated. Although social isolation did not significantly increase this behavior comparing knockouts group housed to social isolated, the greater increase in anxiety observed for *adgrl3.1*^*-/-*^ social isolated raises questions on how social isolation can negatively impact comorbidities associated with EDs. Social isolation has been shown to predict increased comorbid anxiety disorders in some EDs (Butler et al., 2016); for example, anxiety is the most common ADHD comorbidity (Quenneville et al., 2022). In this study, we showed, for the first time, that this internalizing behavior is increased in *adgrl3.1* knockouts. Considering that social isolation tends to decrease anxiety level in WT fish, this shows a gene and environment interaction, where mutants for the *adgrl3.1* show a opposite trend in terms of anxiety, after exposure to social isolation. However, externalizing behaviors such as risk-taking was not affected by social isolation, indicating that impulsive-related phenotypes may not be affected by this environmental factor.

In humans, social isolation is known to alter memory and cognition in healthy individuals where loneliness for 4 years shows a decrease in cognitive function compared to those that had a better social environment (Cacioppo and Cacioppo, 2014). In zebrafish, the effects of social isolation on social and anxiety-like behavior are well-known (Giacomini et al., 2015; Parker et al., 2012; Shams et al., 2018; Shams et al., 2015); however, the effects of the 14 days isolation protocol in adult memory and cognition are still under-explored. Although we did not find any differences in working memory or repetitive behavior, WT animals exposed to social isolation had a decline in their cognitive flexibility by not reaching a peak in their alternations at 40 min and showed decreased exploration. A peak at 40 min in alternations is a normal pattern observed in WT fish in normal housing conditions and is associated with high cognitive flexibility (Cleal et al., 2021). Interestingly, like the anxiety-related findings, social isolation had a greater negative impact on memory and cognition for socially isolated *adgrl3.1* zebrafish, and this was the only group to show decreased working memory compared to WT group-housed. Both *adgrl3.1*^*-/-*^ groups also showed increased repetitive behavior, with greater effects for socially isolated animals. The presence of abnormal repetitive behaviors (ARBs) is a strategy often observed in mice (Langen et al., 2011), primates (Pomerantz et al., 2012) and humans (Manor-Binyamini and Schreiber-Divon 2019) as a coping mechanism when individuals are stressed.

The mechanisms underlying the presence of ARBs may involve the DA system, since exposure to the D1 antagonist SCH-23390 increases the number of repetitions (Cleal et al, 2020) and pretreatment with a DRD1/DRD5 agonist prevents ARB induced by an acute stressor in adult zebrafish (Fontana et al., 2021). Is also known that acute (24-h) and chronic (6-month) isolation decrease the brain levels of dopamine, DOPAC and 5-HIAA in adult zebrafish, together with lower whole-body cortisol levels in chronically isolated animals (Shams et al., 2017). Although the mechanisms underlying social isolation increasing ARB in *adgrl3.1* knockout are yet not clear, transient knockdown of *adgrl3.1* showed altered sensitivity to stimulation of dopamine signaling at larval stages (Lange et al., 2018). Thus, it could be that the DA system is involved in the increased ARBs showed by *adgrl3.1*^*-/-*^.

Social behaviors were also differently affected depending on gene and environment. When looking at *adgrl3.1* knockout, social isolation had an opposite tendency compared to WT, where *adgrl3.1* group-housed show reduced shoaling and socially isolated *adgrl3.1* animals have no significant difference compared to all groups. Impaired social function has been widely reported in ADHD patients as a result of patients being unable to maintain a focus of attention on information within working memory while simultaneously dividing attention among multiple, ongoing events and social cues occurring within the environment (Kofler et al., 2011). It is known that *adgrl3.1* knockout zebrafish exhibit an inattentive behavioral phenotype (Fontana et al., 2023), thus, the deficits in social behavior observed here (higher shoal area and distance inter-fish) could result from the animals’ inability to divide attention among social cues when exposed to a new environment in the shoaling test. Interestingly, social deprivation is known to evoke midbrain craving similar to hunger in humans, supporting the hypothesis that acute isolation causes social craving (Tomova et al., 2020). After two weeks of social isolation *adgrl3.1* may have increased attention focus on social cues due lack of social interaction or increased levels of defensive behavior. As a result, *adgrl3.1* knockout submitted to social isolation showed a strong gene-environment interaction, where social isolation decreased social deficits showed by *adgrl3.1* knockout.

In our previous work, we found that, in addition to displaying core ED behaviors, molecular analysis of *adgrl3.1* knockout zebrafish revealed enriched clusters of differentially expressed genes (DEGs) related to immune signaling (e.g., NACHT, Leucine-rich repeat, P-loop containing nucleoside triphosphatase hydrolase) and transcriptional regulation (e.g., Zinc finger, RNA polymerase II transcription factor activity). These pathways are crucial for intracellular signal transduction, stress response, and immune regulation (Fontana et al., 2023). In a separate study (Alnassar et al., 2023), we found that social isolation of adult zebrafish resulted in significant changes in gene expression, particularly in genes involved in immune responses and structural integrity, such as *SPINT1, COL1A1, COL1A2*, and *S100A4/A5*. These genes overlap significantly with the enriched clusters observed in *adgrl3.1* knockout zebrafish, particularly those related to immune function (e.g., Interleukin [IL]-1 signaling) and structural components (e.g., collagen genes). The convergence of our previous findings strongly suggests that social isolation may exacerbate the phenotypic expression of *adgrl3.1* dysfunction. For example, the role of IL-1 signaling in stress responses and social behavior (DiSabato et al., 2021; Goshen and Yirmiya, 2009), coupled with the identification of immune and structural genes in both studies, supports the hypothesis that social isolation could amplify the behavioral and molecular abnormalities associated with *adgrl3.1* knockout. This interaction is particularly relevant given the inflammatory responses observed in both contexts, indicating that social isolation may have heightened the inflammatory state (Koyama et al., 2021), thereby worsening the *adgrl3.1*-related phenotypes.

## 5. Conclusion

Overall, our findings demonstrate that social isolation has a differential impact on zebrafish behavior based on genotype. Zebrafish knockout for *adgrl3.1* were shown for the first time, to not only exhibit externalizing behaviors but also internalizing behaviors such as increased anxiety and ARBs. When these mutants were socially isolated, an opposite effect was observed compared to WT animals in a variety of complex behaviors, including working memory, anxiety, ARBs and shoaling. Collectively, these data suggest that social isolation differently affects behavior depending on genotype. In addition, social isolation negatively impacts *adgrl3.1* knockout, which could aggravate the negative symptoms and comorbidities associated with EDs. The two-week social isolation paradigm proved to be an effective method for assessing environmental impacts on EDs, highlighting the utility of zebrafish as a translational model. Given the significant overlap of immune-related gene expression changes previously observed in socially isolated WT and *adgrl3.1* knockout zebrafish, future studies should explore the role of the inflammatory system in modulating these behavioral and molecular phenotypes.

## Acknowledgments

The authors would like to thank the School of Pharmacy and Biomedical Sciences, University of Portsmouth, UK, for providing the facilities to conduct the experiments. This study was financed in part by the Coordenação de Aperfeiçoamento de Pessoal de Nível Superior - Brazil (CAPES) - Finance Code 001. MOP received funding from NC3Rs (UK). The funders had no role in study design, data collection, and analysis, decision to publish, or preparation of the manuscript.

## References

Alnassar, N., Hillman, C., Fontana, B.D., Robson, S.C., Norton, W.H.J., Parker, M.O., 2023. angptl4 gene expression as a marker of adaptive homeostatic response to social isolation across the lifespan in zebrafish. Neurobiol Aging 131, 209–221.

Arcos-Burgos, M., Jain, M., Acosta, M.T., Shively, S., Stanescu, H., Wallis, D., Domene, S., Velez, J.I., Karkera, J.D., Balog, J., Berg, K., Kleta, R., Gahl, W.A., Roessler, E., Long, R., Lie, J., Pineda, D., Londono, A.C., Palacio, J.D., Arbelaez, A., Lopera, F., Elia, J., Hakonarson, H., Johansson, S., Knappskog, P.M., Haavik, J., Ribases, M., Cormand, B., Bayes, M., Casas, M., Ramos-Quiroga, J.A., Hervas, A., Maher, B.S., Faraone, S.V., Seitz, C., Freitag, C.M., Palmason, H., Meyer, J., Romanos, M., Walitza, S., Hemminger, U., Warnke, A., Romanos, J., Renner, T., Jacob, C., Lesch, K.P., Swanson, J., Vortmeyer, A., Bailey-Wilson, J.E., Castellanos, F.X., Muenke, M., 2010. A common variant of the latrophilin 3 gene, LPHN3, confers susceptibility to ADHD and predicts effectiveness of stimulant medication. Mol Psychiatry 15(11), 1053–1066.

Butler, T.R., Karkhanis, A.N., Jones, S.R., Weiner, J.L., 2016. Adolescent Social Isolation as a Model of Heightened Vulnerability to Comorbid Alcoholism and Anxiety Disorders. Alcoholism: Clinical and Experimental Research 40(6), 1202–1214.

Cacioppo, J.T., Cacioppo, S., 2014. Older adults reporting social isolation or loneliness show poorer cognitive function 4 years later. 17(2), 59–60.

Cacioppo, J.T., Ernst, J.M., Burleson, M.H., McClintock, M.K., Malarkey, W.B., Hawkley, L.C., Kowalewski, R.B., Paulsen, A., Hobson, J.A., Hugdahl, K., Spiegel, D., Berntson, G.G., 2000. Lonely traits and concomitant physiological processes: the MacArthur social neuroscience studies. Int J Psychophysiol 35(2-3), 143–154.

Canzian, J., Fontana, B.D., Quadros, V.A., Rosemberg, D.B., 2017. Conspecific alarm substance differently alters group behavior of zebrafish populations: Putative involvement of cholinergic and purinergic signaling in anxiety- and fear-like responses. Behav Brain Res 320, 255–263.

Carpenter Rich, E., Loo, S.K., Yang, M., Dang, J., Smalley, S.L., 2009. Social functioning difficulties in ADHD: association with PDD risk. Clin Child Psychol Psychiatry 14(3), 329–344.

Cleal, M., Fontana, B.D., Ranson, D.C., McBride, S.D., Swinny, J.D., Redhead, E.S., Parker, M.O., 2021. The Free-movement pattern Y-maze: A cross-species measure of working memory and executive function. Behav Res Methods(53), 536 – 557.

Cleal, M., Parker, M.O., 2018. Moderate developmental alcohol exposure reduces repetitive alternation in a zebrafish model of fetal alcohol spectrum disorders. Neurotoxicol Teratol 70, 1–9.

de Lacy, N., Ramshaw, M.J., 2023. Selectively predicting the onset of ADHD, oppositional defiant disorder, and conduct disorder in early adolescence with high accuracy. Front Psychiatry 14, 1280326.

DiSabato, D.J., Nemeth, D.P., Liu, X., Witcher, K.G., O’Neil, S.M., Oliver, B., Bray, C.E., Sheridan, J.F., Godbout, J.P., Quan, N., 2021. Interleukin-1 receptor on hippocampal neurons drives social withdrawal and cognitive deficits after chronic social stress. Molecular Psychiatry 26(9), 4770–4782.

Domene, S., Stanescu, H., Wallis, D., Tinloy, B., Pineda, D.E., Kleta, R., Arcos-Burgos, M., Roessler, E., Muenke, M., 2011. Screening of human LPHN3 for variants with a potential impact on ADHD susceptibility. Am J Med Genet B Neuropsychiatr Genet 156B(1), 11-18.

Egan, R.J., Bergner, C.L., Hart, P.C., Cachat, J.M., Canavello, P.R., Elegante, M.F., Elkhayat, S.I., Bartels, B.K., Tien, A.K., Tien, D.H., Mohnot, S., Beeson, E., Glasgow, E., Amri, H., Zukowska, Z., Kalueff, A.V., 2009. Understanding behavioral and physiological phenotypes of stress and anxiety in zebrafish. Behav Brain Res 205(1), 38–44.

Fabricius, K., Helboe, L., Steiniger-Brach, B., Fink-Jensen, A., Pakkenberg, B., 2010. Stereological brain volume changes in post-weaned socially isolated rats. Brain Res 1345, 233–239.

Fontana, B.D., Cleal, M., Clay, J.M., Parker, M.O., 2019. Zebrafish (Danio rerio) behavioral laterality predicts increased short-term avoidance memory but not stress-reactivity responses. Animal cognition.

Fontana, B.D., Cleal, M., Gibbon, A.J., McBride, S.D., Parker, M.O., 2021. The effects of two stressors on working memory and cognitive flexibility in zebrafish (Danio rerio): The protective role of D1/D5 agonist on stress responses. Neuropharmacology 196, 108681.

Fontana, B.D., Cleal, M., Parker, M.O., 2020. Female adult zebrafish (Danio rerio) show higher levels of anxiety-like behavior than males, but do not differ in learning and memory capacity. Eur J Neurosci 52(1), 2604–2613.

Fontana, B.D., Reichmann, F., Tilley, C.A., Lavlou, P., Shkumatava, A., Alnassar, N., Hillman, C., Karlsson, K.Ö., Norton, W.H.J., Parker, M.O., 2023. adgrl3.1-deficient zebrafish show noradrenaline-mediated externalizing behaviors, and altered expression of externalizing disorder-candidate genes, suggesting functional targets for treatment. Translational Psychiatry 13(1), 304.

Forsatkar, M.N., Safari, O., Boiti, C., 2017. Effects of social isolation on growth, stress response, and immunity of zebrafish. acta ethologica 20(3), 255–261.

Giacomini, A.C., de Abreu, M.S., Koakoski, G., Idalencio, R., Kalichak, F., Oliveira, T.A., da Rosa, J.G., Gusso, D., Piato, A.L., Barcellos, L.J., 2015. My stress, our stress: blunted cortisol response to stress in isolated housed zebrafish. Physiol Behav 139, 182–187.

Goshen, I., Yirmiya, R., 2009. Interleukin-1 (IL-1): A central regulator of stress responses. Frontiers in Neuroendocrinology 30(1), 30–45.

Green, J., Collins, C., Kyzar, E.J., Pham, M., Roth, A., Gaikwad, S., Cachat, J., Stewart, A.M., Landsman, S., Grieco, F., Tegelenbosch, R., Noldus, L.P., Kalueff, A.V., 2012. Automated high-throughput neurophenotyping of zebrafish social behavior. J Neurosci Methods 210(2), 266–271.

Gross, A.N., Engel, A.K., Richter, S.H., Garner, J.P., Wurbel, H., 2011. Cage-induced stereotypies in female ICR CD-1 mice do not correlate with recurrent perseveration. Behav Brain Res 216(2), 613–620.

Hoza, B., 2007. Peer Functioning in Children With ADHD. Journal of Pediatric Psychology 32(6), 655–663.

Hwang, I.W., Lim, M.H., Kwon, H.J., Jin, H.J., 2015. Association of LPHN3 rs6551665 A/G polymorphism with attention deficit and hyperactivity disorder in Korean children. Gene 566(1), 68–73.

Ieraci, A., Mallei, A., Popoli, M., 2016. Social Isolation Stress Induces Anxious-Depressive-Like Behavior and Alterations of Neuroplasticity-Related Genes in Adult Male Mice. Neural Plasticity 2016, 6212983.

Kalueff, A.V., Gebhardt, M., Stewart, A.M., Cachat, J.M., Brimmer, M., Chawla, J.S., Craddock, C., Kyzar, E.J., Roth, A., Landsman, S., Gaikwad, S., Robinson, K., Baatrup, E., Tierney, K., Shamchuk, A., Norton, W., Miller, N., Nicolson, T., Braubach, O., Gilman, C.P., Pittman, J., Rosemberg, D.B., Gerlai, R., Echevarria, D., Lamb, E., Neuhauss, S.C., Weng, W., Bally-Cuif, L., Schneider, H., Zebrafish Neuroscience Research, C., 2013. Towards a comprehensive catalog of zebrafish behavior 1.0 and beyond. Zebrafish 10(1), 70–86.

Kofler, M.J., Rapport, M.D., Bolden, J., Sarver, D.E., Raiker, J.S., Alderson, R.M., 2011. Working memory deficits and social problems in children with ADHD. J Abnorm Child Psychol 39(6), 805–817.

Koyama, Y., Nawa, N., Yamaoka, Y., Nishimura, H., Sonoda, S., Kuramochi, J., Miyazaki, Y., Fujiwara, T., 2021. Interplay between social isolation and loneliness and chronic systemic inflammation during the COVID-19 pandemic in Japan: Results from U-CORONA study. Brain Behav Immun 94, 51–59.

Kwan, C., Gitimoghaddam, M., Collet, J.P., 2020. Effects of Social Isolation and Loneliness in Children with Neurodevelopmental Disabilities: A Scoping Review. Brain Sci 10(11).

Lange, M., Froc, C., Grunwald, H., Norton, W.H.J., Bally-Cuif, L., 2018. Pharmacological analysis of zebrafish lphn3.1 morphant larvae suggests that saturated dopaminergic signaling could underlie the ADHD-like locomotor hyperactivity. Progress in Neuro-Psychopharmacology and Biological Psychiatry 84, 181–189.

Lange, M., Norton, W., Coolen, M., Chaminade, M., Merker, S., Proft, F., Schmitt, A., Vernier, P., Lesch, K.P., Bally-Cuif, L., 2012. The ADHD-susceptibility gene lphn3.1 modulates dopaminergic neuron formation and locomotor activity during zebrafish development. Molecular Psychiatry 17(9), 946–954.

Langen, M., Kas, M.J., Staal, W.G., van Engeland, H., Durston, S., 2011. The neurobiology of repetitive behavior: of mice. Neurosci Biobehav Rev 35(3), 345–355.

Lapiz, M.D., Fulford, A., Muchimapura, S., Mason, R., Parker, T., Marsden, C.A., 2003. Influence of postweaning social isolation in the rat on brain development, conditioned behavior, and neurotransmission. Neurosci Behav Physiol 33(1), 13–29.

Levin, E.D., Bencan, Z., Cerutti, D.T., 2007. Anxiolytic effects of nicotine in zebrafish. Physiol Behav 90(1), 54–58.

Lilienfeld, S.O., 2003. Comorbidity between and within childhood externalizing and internalizing disorders: reflections and directions. J Abnorm Child Psychol 31(3), 285–291.

Liu, J., 2004. Childhood externalizing behavior: theory and implications. J Child Adolesc Psychiatr Nurs 17(3), 93–103.

Manor-Binyamini, I., Schreiber-Divon, M., 2019. Repetitive behaviors: Listening to the voice of people with high-functioning autism spectrum disorder. Research in Autism Spectrum Disorders 64, 23–30.

Matthews, G.A., Tye, K.M., 2019. Neural mechanisms of social homeostasis. Ann N Y Acad Sci 1457(1), 5–25.

McMillan, D.R., Kayes-Wandover, K.M., Richardson, J.A., White, P.C., 2002. Very large G protein-coupled receptor-1, the largest known cell surface protein, is highly expressed in the developing central nervous system. J Biol Chem 277(1), 785–792.

Meßler, C.F., Holmberg, H.-C., Sperlich, B., 2016. Multimodal Therapy Involving High-Intensity Interval Training Improves the Physical Fitness, Motor Skills, Social Behavior, and Quality of Life of Boys With ADHD: A Randomized Controlled Study. Journal of Attention Disorders 22(8), 806–812.

Muller, T.E., Nunes, S.Z., Silveira, A., Loro, V.L., Rosemberg, D.B., 2017. Repeated ethanol exposure alters social behavior and oxidative stress parameters of zebrafish. Prog Neuropsychopharmacol Biol Psychiatry 79(Pt B), 105–111.

Mumtaz, F., Khan, M.I., Zubair, M., Dehpour, A.R., 2018. Neurobiology and consequences of social isolation stress in animal model-A comprehensive review. Biomed Pharmacother 105, 1205–1222.

Norton, W.H., Stumpenhorst, K., Faus-Kessler, T., Folchert, A., Rohner, N., Harris, M.P., Callebert, J., Bally-Cuif, L., 2011. Modulation of Fgfr1a signaling in zebrafish reveals a genetic basis for the aggression-boldness syndrome. J Neurosci 31(39), 13796–13807.

Parker, M.O., Millington, M.E., Combe, F.J., Brennan, C.H., 2012. Housing conditions differentially affect physiological and behavioural stress responses of zebrafish, as well as the response to anxiolytics. PLoS One 7(4), e34992.

Pomerantz, O., Paukner, A., Terkel, J., 2012. Some stereotypic behaviors in rhesus macaques (Macaca mulatta) are correlated with both perseveration and the ability to cope with acute stressors. Behav Brain Res 230(1), 274–280.

Quenneville, A.F., Kalogeropoulou, E., Nicastro, R., Weibel, S., Chanut, F., Perroud, N., 2022. Anxiety disorders in adult ADHD: A frequent comorbidity and a risk factor for externalizing problems. Psychiatry Research 310, 114423.

Ray, A.R., Evans, S.W., Langberg, J.M., 2017. Factors Associated with Healthy and Impaired Social Functioning in Young Adolescents with ADHD. J Abnorm Child Psychol 45(5), 883–897.

Rosemberg, D.B., Braga, M.M., Rico, E.P., Loss, C.M., Cordova, S.D., Mussulini, B.H., Blaser, R.E., Leite, C.E., Campos, M.M., Dias, R.D., Calcagnotto, M.E., de Oliveira, D.L., Souza, D.O., 2012. Behavioral effects of taurine pretreatment in zebrafish acutely exposed to ethanol. Neuropharmacology 63(4), 613–623.

Samek, D.R., Hicks, B.M., 2014. Externalizing Disorders and Environmental Risk: Mechanisms of Gene-Environment Interplay and Strategies for Intervention. Clin Pract (Lond) 11(5), 537–547.

Schmidel, A.J., Assmann, K.L., Werlang, C.C., Bertoncello, K.T., Francescon, F., Rambo, C.L., Beltrame, G.M., Calegari, D., Batista, C.B., Blaser, R.E., Roman Junior, W.A., Conterato, G.M., Piato, A.L., Zanatta, L., Magro, J.D., Rosemberg, D.B., 2014. Subchronic atrazine exposure changes defensive behaviour profile and disrupts brain acetylcholinesterase activity of zebrafish. Neurotoxicol Teratol 44, 62–69.

Schoenfeld, T.J., Gould, E., 2012. Stress, stress hormones, and adult neurogenesis. Exp Neurol 233(1), 12–21.

Scholz, N., Gehring, J., Guan, C., Ljaschenko, D., Fischer, R., Lakshmanan, V., Kittel, R.J., Langenhan, T., 2015. The adhesion GPCR latrophilin/CIRL shapes mechanosensation. Cell Rep 11(6), 866–874.

Serra, M., Pisu, M.G., Floris, I., Biggio, G., 2005. Social isolation-induced changes in the hypothalamic-pituitary-adrenal axis in the rat. Stress 8(4), 259–264.

Shams, S., Amlani, S., Buske, C., Chatterjee, D., Gerlai, R., 2018. Developmental social isolation affects adult behavior, social interaction, and dopamine metabolite levels in zebrafish. Dev Psychobiol 60(1), 43–56.

Shams, S., Chatterjee, D., Gerlai, R., 2015. Chronic social isolation affects thigmotaxis and whole-brain serotonin levels in adult zebrafish. Behav Brain Res 292, 283–287.

Shams, S., Seguin, D., Facciol, A., Chatterjee, D., Gerlai, R., 2017. Effect of social isolation on anxiety-related behaviors, cortisol, and monoamines in adult zebrafish. Behav Neurosci 131(6), 492–504.

Shaw-Zirt, B., Popali-Lehane, L., Chaplin, W., Bergman, A., 2005. Adjustment, Social Skills, and Self-Esteem in College Students With Symptoms of ADHD. Journal of Attention Disorders 8(3), 109–120.

Sih, A., Bell, A.M., Johnson, J.C., Ziemba, R.E., 2004. Behavioral syndromes: an intergrative overiew. Q Rev Biol 79(3), 241–277.

Smit, S., Mikami, A.Y., Normand, S., 2020. Correlates of Loneliness in Children with Attention-Deficit/Hyperactivity Disorder: Comorbidities and Peer Problems. Child Psychiatry Hum Dev 51(3), 478–489.

Speyer, L.G., Eisner, M., Ribeaud, D., Luciano, M., Auyeung, B., Murray, A.L., 2021. Developmental Relations Between Internalising Problems and ADHD in Childhood: a Symptom Level Perspective. Res Child Adolesc Psychopathol 49(12), 1567–1579.

Sveinsdóttir, H.S., Christensen, C., Þorsteinsson, H., Lavalou, P., Parker, M.O., Shkumatava, A., Norton, W.H.J., Andriambeloson, E., Wagner, S., Karlsson, K.Ö., 2023. Novel non-stimulants rescue hyperactive phenotype in an adgrl3.1 mutant zebrafish model of ADHD. Neuropsychopharmacology 48(8), 1155–1163.

Tomova, L., Wang, K.L., Thompson, T., Matthews, G.A., Takahashi, A., Tye, K.M., Saxe, R., 2020. Acute social isolation evokes midbrain craving responses similar to hunger. Nature neuroscience 23(12), 1597–1605.

Tuchscherer, M., Kanitz, E., Puppe, B., Tuchscherer, A., Stabenow, B., 2004. Effects of postnatal social isolation on hormonal and immune responses of pigs to an acute endotoxin challenge. Physiology & Behavior 82(2), 503–511.

Westenbroek, C., Den Boer, J.A., Veenhuis, M., Ter Horst, G.J., 2004. Chronic stress and social housing differentially affect neurogenesis in male and female rats. Brain Res Bull 64(4), 303–308.

Wong, K., Elegante, M., Bartels, B., Elkhayat, S., Tien, D., Roy, S., Goodspeed, J., Suciu, C., Tan, J., Grimes, C., Chung, A., Rosenberg, M., Gaikwad, S., Denmark, A., Jackson, A., Kadri, F., Chung, K.M., Stewart, A., Gilder, T., Beeson, E., Zapolsky, I., Wu, N., Cachat, J., Kalueff, A.V., 2010. Analyzing habituation responses to novelty in zebrafish (Danio rerio). Behav Brain Res 208(2), 450–457.

Wright, D., Rimmer, L.B., Pritchard, V.L., Krause, J., Butlin, R.K., 2003. Inter and intra-population variation in shoaling and boldness in the zebrafish (Danio rerio). Naturwissenschaften 90(8), 374–377.

Zlatković, J., Todorović, N., Bošković, M., Pajović, S.B., Demajo, M., Filipović, D., 2014. Different susceptibility of prefrontal cortex and hippocampus to oxidative stress following chronic social isolation stress. Molecular and Cellular Biochemistry 393(1), 43–57.

